## Supplementary material for "Impaired motor-to-sensory transformation mediates auditory hallucinations" (from another version of this preprint: version 2, 2024-02-03)

*Corresponding Author.


**This PDF file includes:**

**Figures S1 to S4**

**Table S1**

**Figure S1. The modulation effects on auditory responses during general preparation (GP) in normal control group**

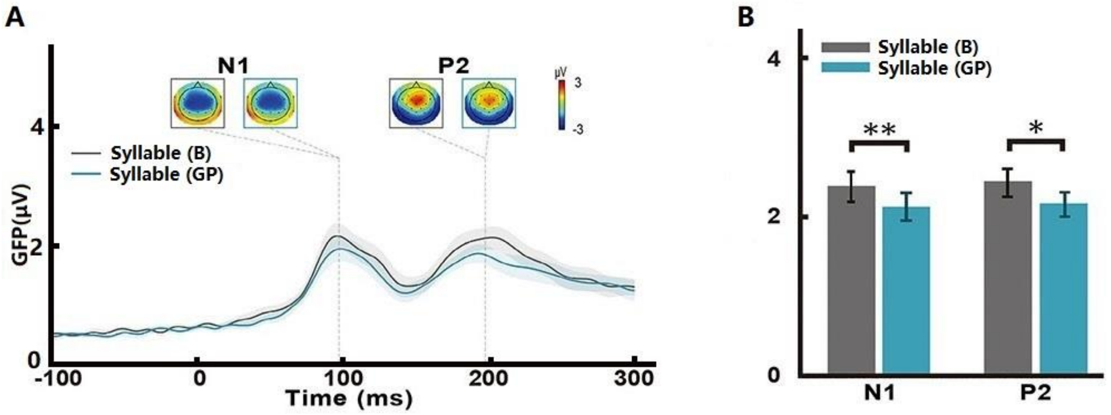


*Figure S1* (A) ERP time course and topographic responses for GP and B conditions in normal populations. Typical N1 and P2 components were observed in the GFP waveforms for each condition. The response topographies at each peak latency are shown in boxes with the same color code of conditions. (B) Mean GFP amplitude at N1 and P2 latencies for GP (blue) and B (gray) conditions in normal populations.

In normal population, the amplitude of early N1 response in GP was less than that in B ($t(18) = 3.406$, $p = 0.003$, $d = 0.3$). The amplitude of later P2 response in GP was reduced relative to B ($t(18) = 2.240$, $p = 0.038$, $d = 0.342$) (We are requiring permission for using the normals' results from the original publisher). These results suggested that CD that was induced during GP suppressed auditory responses.

**Figure S2. The modulation effects on auditory responses during specific preparation(SP) in normal control group**

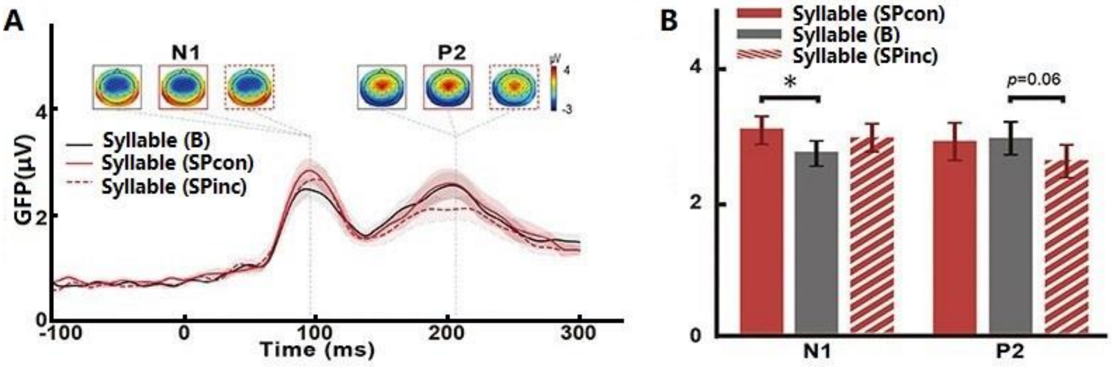


*Figure S2* (A) ERP time course and topographic responses for SP and B conditions in non-AVHs patients. (B) Mean GFP amplitudes at N1 and P2 latencies for SP and B conditions. Responses in SPcon were significantly larger than those in B in N1 components

In the SP, early neural responses of N1 were larger than that in B when the auditory syllables were congruent with the SP visual cues (SPcon) ($t(15) = -2.49$, $p = 0.025$, $d = -0.432$). The effect was not significant in the later auditory responses of P2 ($t(15) = 0.248$, $p = 0.808$, $d = 0.039$). However, when the auditory syllables were incongruent with the specific preparation (SPinc), the effect in N1was not significant ($t(15) = -1.48$, $p = 0.160$, $d = -0.283$), nor in P2 ($t(15) = 2.024$, $p = 0.061$, $d = 0.342$) (We are requiring permission for using the normals' results from the original publisher). These results suggested that motor signals during SP modulated the perceptual responses based on the content congruency.

**Figure S3. The absence of modulation effects on tones during general preparation**

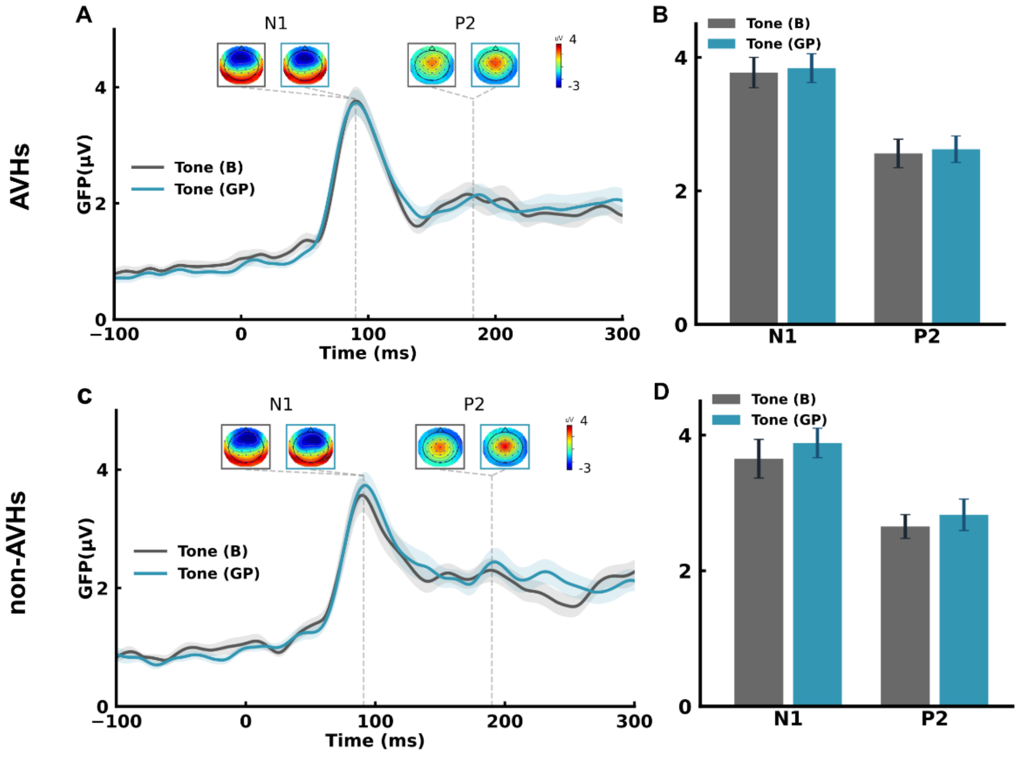


*Figure S3* (A) ERP time course and topographic responses for GP and B conditions in AVHs patients. Typical N1 and P2 components were observed in the GFP waveforms for each condition. The response topographies at each peak latency are shown in boxes with the same color code of conditions. (B) Mean GFP amplitude at N1 and P2 latencies for GP (blue) and B (gray) conditions in AVHs patients. (C) ERP time course and topographic responses for GP and B conditions in non-AVHs patients. (D) Mean GFP amplitudes at the N1 and P2 latencies for GP (blue) and B (gray) conditions in non-AVHs patients. No significant differences between GP and B were observed in either group. Error bars indicate ± SEMs.

**Figure S4. The absence of modulation effects on tones during specific preparation**

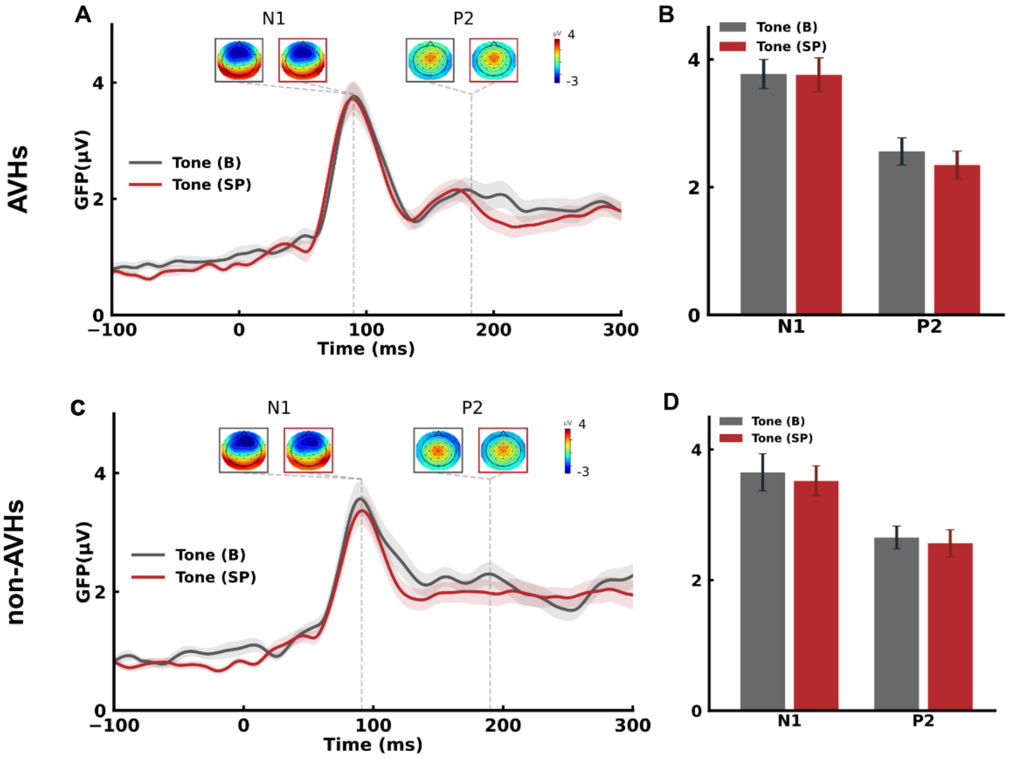


*Figure S4* (A) ERP time course and topographic responses for SP and B conditions in AVHs patients. Typical N1 and P2 components were observed in the GFP waveforms of each condition. The response topographies at each peak latency are shown in boxes with the same color code of conditions. (B) Mean GFP amplitude at N1 and P2 latencies for SP (red) and B (gray) conditions in AVHs patients. (C) ERP time course and topographic responses for SP and B conditions in non-AVHs patients. (D) Mean GFP amplitudes at the N1 and P2 latencies for SP (red) and B (gray) conditions in non-AVHs patients. No significant differences between SP and B were observed in either group. Error bars indicate ± SEMs.

**Table S1. Demographics of AVHs, non-AVHs, and Normal Controls**

| | AVHs (n=20) | non-AVHs (n=20) | Normal Controls for GP (n=19) | F or $\chi^2$ | *p* | Normal Controls for SP (n=16) | F or $\chi^2$ | *p* |
|---|---|---|---|---|---|---|---|---|
| **Gender (M/F)** | 10/10 | 14/6 | 5/14 | 7.45 | 0.024 | 5/11 | 5.39 | 0.07 |
| **Age (years)** | 25.60 (6.75) | 30.75 (6.15) | 23.89 (2.96) | 8.06 | 0.001 | 23.13 (3.38) | 8.40 | 0.001 |
| **Education (years)** | 13.00 (2.27) | 14.07 (3.16) | 16.53 (2.50) | 8.87 | 0.001 | 15.81 (2.79) | 4.64 | 0.014 |
| **Age of onset** | 20.85 (6.51) | 25.20 (7.70) | N/A | 1.93 | 0.06 | N/A | N/A | N/A |
| **Duration (month)** | 68.50 (65.22) | 66.00 (46.27) | N/A | 0.14 | 0.89 | N/A | N/A | N/A |
| **PANSS total score** | 76.50 (15.58) | 67.75 (13.91) | N/A | 1.87 | 0.07 | N/A | N/A | N/A |
| **P3 subscore** | 4.35 (1.27) | 1.00 (0.0) | N/A | 11.82 | <0.001 | N/A | N/A | N/A |
| **P subscore** | 21.25 (5.39) | 17.10 (4.06) | N/A | 2.75 | 0.009 | N/A | N/A | N/A |
| **N subscore** | 18.50 (4.47) | 17.90 (4.48) | N/A | 0.42 | 0.67 | N/A | N/A | N/A |
| **G subscore** | 36.75 (8.78) | 32.75 (8.08) | N/A | 1.50 | 0.14 | N/A | N/A | N/A |
| **AVHs total score** | 26.15 (4.42) | N/A | N/A | N/A | N/A | N/A | N/A | N/A |

Notes: PANSS, Positive and Negative Syndrome Scale; P subscore, positive symptom subscore; N subscore, negative symptom subscore; G subscore, general psychopathology subscore; N/A, not applicable. The standard deviation is in parentheses.
